# Ferroptosis Regulation by the NGLY1/NFE2L1 Pathway

**DOI:** 10.1101/2021.10.12.463965

**Authors:** Giovanni C. Forcina, Lauren Pope, Magdalena Murray, Wentao Dong, Monther Abu-Remaileh, Carolyn R. Bertozzi, Scott J. Dixon

**Affiliations:** Department of Biology, Stanford University, Stanford CA 94305, USA; Department of Chemical Engineering & Stanford ChEM-H, Stanford University, Stanford CA 94305, USA; Department of Chemistry & Stanford ChEM-H, Stanford University, Stanford, CA 94305; Howard Hughes Medical Institute, Stanford, CA 94305

**Keywords:** NGLY1, NFE2L1, ferroptosis, CNC transcription factors

## Abstract

Ferroptosis is an oxidative form of non-apoptotic cell death whose transcriptional regulation is poorly understood. Cap’n’collar (CNC) transcription factors including Nuclear Factor Erythroid-2 Related Factor 1 (NFE2L1/NRF1) and NFE2L2 (NRF2) are important regulators of oxidative stress responses. Here, we report that NFE2L1 expression inhibits ferroptosis, independent of NFE2L2. NFE2L1 inhibits ferroptosis by promoting expression of the key anti-ferroptotic lipid hydroperoxidase glutathione peroxidase 4 (GPX4). NFE2L1 abundance and function are regulated post-translationally by N-glycosylation. Functional maturation of NFE2L1 requires deglycosylation by cytosolic peptide:N-glycanase 1 (NGLY1). We find that loss of NGLY1 or NFE2L1 enhances ferroptosis sensitivity. Expression of wild-type NGLY1 but not a disease-associated NGLY1 mutant inhibits ferroptosis, and this effect is dependent on the presence of NFE2L1. Enhanced ferroptosis sensitivity in NFE2L1 and NFE2L2 knockout cells can be potently reverted by expression of an NFE2L1 mutant containing eight asparagine-to-aspartate protein sequence substitutions, which mimic NGLY1-catalyzed sequence editing. Enhanced ferroptosis sensitivity in NGLY1/NFE2L1 pathway mutants could also be reversed by overexpression of NFE2L2. These results suggest that ferroptosis sensitivity is regulated by NGLY1-catalyzed NFE2L1 deglycosylation, and highlight a broad role for CNC transcription factors in ferroptosis regulation.

**Significance Statement:** Ferroptosis is an oxidative form of cell death whose biochemical regulation remains incompletely understood. NFE2L1/NRF1 is a cap’n’collar (CNC) transcription factor whose role in ferroptosis regulation is unclear. Unlike the CNC family member NFE2L2/NRF2, NFE2L1 is an N-glycoprotein whose abundance is regulated by post-translational deglycosylation catalyzed by the enzyme peptide:N-glycanase 1 (NGLY1). Our results indicate that NGLY1-mediated NFE2L1 deglycosylation, resulting in ‘editing’ of the NFE2L1 amino acid sequence, is necessary for NFE2L1 to inhibit ferroptosis. Mechanistically, NFE2L1 inhibits ferroptosis by via the anti-ferroptotic protein GPX4. This work demonstrates that CNC transcription factors beyond NFE2L2 can regulate ferroptosis. This work may suggest a role of misregulation of ferroptosis in NGLY1 deficiency, an ultrarare genetic disorder.

## Introduction

Ferroptosis is an oxidative, non-apoptotic cell death pathway activated in several distinct acute and chronic pathophysiological cell death contexts (1). Ferroptosis can be triggered by depletion of cystine (the oxidized form of cysteine), which is necessary for the synthesis of the anti-ferroptotic metabolites glutathione and coenzyme A, or by direct inactivation of the glutathione-dependent lipid hydroperoxidase GPX4 (2-4). Cystine depletion and/or GPX4 inactivation result in the iron-dependent accumulation of lipid peroxides at the plasma membrane and other sites within the cell, ultimately resulting in membrane permeabilization (2, 5, 6). The factors that modulate ferroptosis sensitivity by impinging on key regulators like GPX4 are incompletely understood. It is of great interest to identify factors that govern ferroptosis sensitivity, as these may suggest novel points of intervention to either enhance or inhibit this process therapeutically.

The cap’n’collar (CNC) family of transcription factors are highly conserved amongst metazoans and play an important role in maintaining homeostasis in response to stress (7). Vertebrates express four CNC family transcription factors, including the related NFE2L1 (NRF1) and NFE2L2 (NRF2) proteins. NFE2L2 is considered the master ‘antioxidant’ transcription factor and can inhibit ferroptosis by regulating the expression of the cystine transporter subunit *SLC7A11*, glutamate-cysteine ligase (GCL) subunits *GCLC* and *GCLM*, and other effectors (8-15). Interestingly, however, *Nfe2l2* is not essential for development in mice, and adult animals exhibit only minor phenotypic abnormalities. By contrast, disruption of *Nfe2l1* is embryonic lethal in mice, suggesting that NFE2L2 and NFE2L1 harbor non-redundant roles (16, 17). Notably, NFE2L1 can also regulate antioxidant gene expression, as well as the expression of various proteasome subunits for which the transcription factor is canonically studied (18-22). Whether NFE2L1 can regulate ferroptosis sensitivity similarly to NFE2L2 is not known.

NFE2L2 and NFE2L1 protein abundance are both regulated post-translationally, but in distinct manners. NFE2L2 is regulated by its cytoplasmic inhibitor Kelch-like ECH-associating protein 1 (KEAP1) (23). Keap1 binds to NFE2L2 in a conserved Neh2 domain also present in NFE2L1, sequesters NFE2L2 in the cytosol, and targets the protein for degradation after ubiquitylation by a Cullin3 (Cul3)-based E3 ubiquitin ligase (24, 25). By contrast, NFE2L1 is constitutively translated into the endoplasmic reticulum, N-glycosylated by oligosaccharyltransferase, translocated from the lumen of the ER into the cytosol, released from the ER membrane into the cytosol by DNA Damage Inducible 1 Homolog 2 (DDI2), and deglycosylated by NGLY1 (26, 27). This deglycosylation reaction leads to ‘editing’ of previously glycosylated Asn residues to Asp residues and is necessary for NFE2L1 to regulate proteasome biogenesis in *C. elegans* (28). Bi-allelic *NGLY1* mutation causes an ultra-rare human disorder called Ngly1 deficiency, characterized by intellectual disability, hyperkinetic movement disorder, liver dysfunction, and alacrima (29). Whether this is due to altered NFE2L1 glycosylation and function is not well understood.

Here, we report that the NGLY1/NFE2L1 pathway regulates ferroptosis sensitivity, in parallel to NFE2L2. Cells lacking NGLY1 or NFE2L1 are sensitized to ferroptosis due to loss of GPX4 expression. NGLY1 is critical for the ability of NFE2L1 to promote GPX4 expression and inhibit ferroptosis via Asn-to-Asp protein sequence editing, which promotes NFE2L1 stability. To this end, we show that a fully ‘edited’ NFE2L1 mutant, NFE2L1^8ND^, is sufficient to inhibit ferroptosis. Moreover, NFE2L1^8ND^ can largely restore ferroptosis sensitivity in cells without NFE2L2. The enhanced ferroptosis sensitivity associated with NGLY1/NFE2L1 pathway disruption can also be compensated for by increased NFE2L2 expression. Thus, related CNC family transcription factors govern ferroptosis sensitivity in parallel through both shared and distinct mechanisms. These results also point to a potential role for misregulated ferroptosis in the ultrarare genetic disorder NGLY1 deficiency.

## Results

### Disruption of CNC Transcription Factors Sensitizes to Ferroptosis

The KEAP1/NFE2L2 and NGLY1/NFE2L1 pathways share the ability to regulate antioxidant gene expression, but the extent to which these pathways may operate independently to regulate oxidative phenotypes like ferroptosis has not been examined. We first analyzed data available from the CTD^2^ consortium, available through the DepMap portal, where it is possible to correlate various molecular readouts and cellular phenotypes across hundreds of cancer cell lines (30, 31). Across over 700 cancer cell lines, we observed a correlation between *KEAP1* mutation status and resistance to erastin-induced or ML162-induced ferroptosis (Fig. 1*A,B*). Thus, the presence of mutant *KEAP1*, which leads to NFE2L2 accumulation, correlates with ferroptosis resistance.

**Figure 1.**
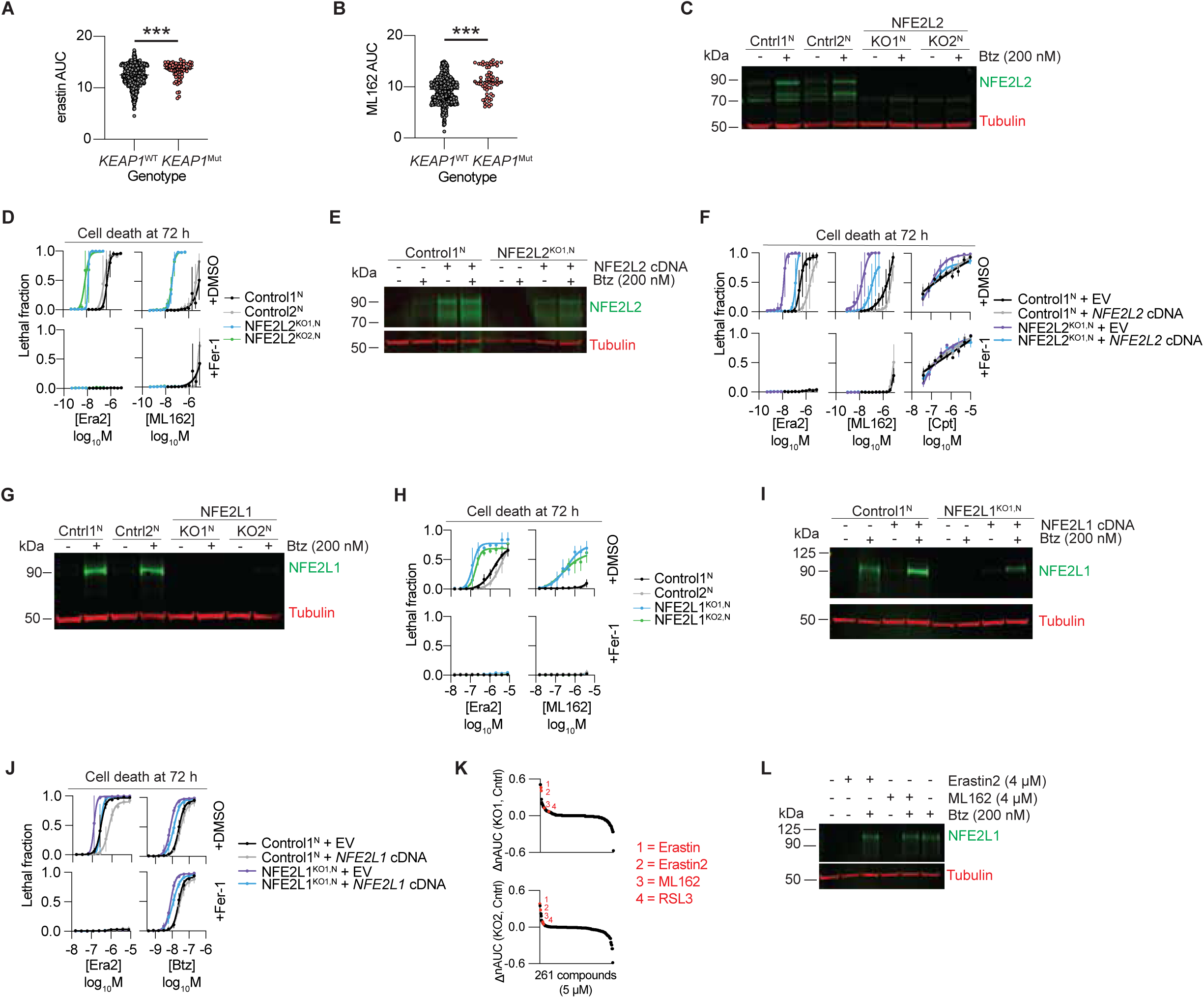
NFE2L2 and NFE2L1 negatively regulate ferroptosis. (*A* and *B*) Correlation between KEAP1 mutation status and sensitivity to ferroptosis inducers erastin2 or ML162. (*C*) NFE2L2 protein levels in Control^N^ and NFE2L2^KO1/2,N^ cell lines. Bortezomib treatment further stabilizes NFE2L2 in Control^N^ cells by blocking proteasomal degradation. (*D*) Ferroptosis sensitivity in Control^N^ and NFE2L2^KO1/2,N^ cell lines treated with a 10-point, 2-fold dose-response of erastin2 and ML162. (*E*) NFE2L2 protein levels in Control1^N^ and NFE2L2^KO1,N^ cells transduced with empty vector (EV) lentivirus or lentivirus directing the expression of NFE2L2 cDNA. (*F*) Ferroptosis sensitivity in Control1^N^ and NFE2L2^KO1,N^ cells transduced with EV or NFE2L2 cDNA. (*G*) NFE2L1 protein levels in Control1/2^N^ and NFE2L1^KO1/2,N^ cell lines. Bortezomib treatment leads to NFE2L1 accumulation in Control1/2^N^ cells. (*H*) Ferroptosis sensitivity in Control1/2^N^ and NFE2L1^KO1/2,N^ cell lines treated with a 10-point, 2-fold dose-response of erastin2 and ML162. (*I*) NFE2L1 protein levels in Control1^N^ and NFE2L1^KO1,N^ cells transduced with EV or NFE2L1 cDNA. (*J*) Ferroptosis sensitivity in Control1^N^ and NFE2L2^KO1,N^ cells overexpressing EV or NFE2L1 cDNA. (*K*) Delta nAUC between NFE2L1^KO1,N^ and Control1^N^, and NFE2L1^KO2,N^ and Control1^N^ after treatment with 261 bioactive small molecules (5 µM). (*L*) Effect of ferroptosis inducers on NFE2L1 protein levels. Cells were treated with vehicle (DMSO), erastin2 (4 µM) or ML162 (4 µM) ± Btz (200 nM) for 4 h. Data in (*D,H*) represent mean ± S.D. of three biological replicates. Data in (*F,J*) represent mean ± S.D. of four biological replicates. Data in (*K*) represent the average difference in normalized area under the curve (ΔnAUC) of three biological replicates. Higher (ΔnAUC) correspond to enhanced cell death for a given treatment in NFE2L1^KO,N^ cells versus Control1^N^ cells. EV, empty vector; Era2, erastin2; Fer-1, ferrostatin-1; Btz, bortezomib; Cpt, camptothecin.

To investigate further, we explored the role of NFE2L2 in human KEAP1-mutant A549 cells, which constitutively accumulate high levels of NFE2L2 (32). We isolated two clonal A549 cell lines where *NFE2L2* was disrupted (i.e. ‘knocked out’, KO) using CRISPR/Cas9 technology, as well as two clonal Control cell lines that underwent the CRISPR process but expressed NFE2L2 normally (Fig. 1*C*). To amplify our ability to detect NFE2L2, we treated cells with bortezomib, which caused the expected accumulation of NFE2L2 in Control1/2 cells but not in NFE2L2^KO1/2^ cells (33) (Fig. 1*C*). To examine cell death, we transduced all four cell lines with lentivirus directing the expression of the live cell marker nuclear-localized mKate2 (denoted by the superscript ‘N’) that, together with incubation with the dead cell dye SYTOX Green, enabled cell death to be directly quantified over time using the scalable analysis of cell death kinetics (STACK) method (34). Compared to Control1/2^N^ cell lines, NFE2L2^KO1/2,N^ cell lines exhibited a remarkable >100-fold heightened sensitivity to two mechanistically distinct ferroptosis-inducing small molecules, the system x_c_^-^ inhibitor erastin2 and the covalent GPX4 inhibitor ML162 (Fig. 1*D*, *SI Appendix*, Table S1 reports all EC_50_ values for all dose-response curves in this work). Cell death under these conditions was confirmed to be ferroptotic because it was fully suppressed by the specific ferroptosis inhibitor ferrostatin-1 (Fer-1) (Fig. 1*D*). The enhanced ferroptosis sensitivity of NFE2L2^KO1,N^ cells was reverted by expression of wild-type *NFE2L2* cDNA, which also enhanced ferroptosis resistance when introduced intro Control1^N^ cells (Fig. 1*E,F*). Thus, constitutively expressed NFE2L2 is an exceptionally potent negative regulator of ferroptosis in A549 cells.

Given the genetic non-redundancy observed between NFE2L2 and NFE2L1 in mammalian development (16, 17), we next examined whether NFE2L1 disruption impacted ferroptosis sensitivity. For these studies we continued to employ A549 cells as a model, as the constitutively high levels of NFE2L2 protein would allow us to ascertain whether NFE2L1 loss resulted in non-redundant phenotypes. We isolated two independent NFE2L1 gene-disrupted clones (Fig. 1*G*). NFE2L1 is constitutively degraded by the proteasome and, in Control1/2^N^ cells, bortezomib treatment resulted in the expected accumulation of NFE2L1 protein (26) (Fig. 1*G*). We transduced these cell lines with mKate2 lentivirus and assayed cell death using STACK. Strikingly, disruption of NFE2L1 significantly enhanced the sensitivity of these cells to erastin2 and ML162 compared to Control1/2^N^ cell lines (Fig. 1*H*). Normal ferroptosis sensitivity was restored to NFE2L1^KO1,N^ cells by re-expression of NFE2L1 (Fig. 1*I,J*). Moreover, NFE2L1 overexpression in Control1^N^ cells further elevated NFE2L1 levels and ferroptosis resistance beyond that achieved by *KEAP1* mutation alone (Fig. 1*I,J*). The sensitization of NFE2L1^KO1/2,N^ cells towards ferroptosis was relatively specific, as they exhibited heightened sensitivity to erastin2, erastin, ML162, a distinct GPX4 inhibitor, RSL3, and only a small number of other compounds from a set of 261 bioactive molecules, compared to Control1^N^ cells (Fig. 1*K*). We conclude that NFE2L1 inhibits ferroptosis independently of NFE2L2.

As noted above, proteasome inhibition causes NFE2L1 accumulation, which then can induce a transcriptional response that promotes proteasome biosynthesis (21, 22, 26). However, NFE2L1 can also regulate the expression of various antioxidant genes (19, 20). We therefore wondered whether pro-ferroptotic oxidative stress could trigger NFE2L1 accumulation, and whether this influenced ferroptosis sensitivity. Unlike bortezomib, neither erastin2 nor ML162 triggered NFE2L1 accumulation alone, and combinations of these agents with bortezomib did not further increase NFE2L1 protein levels (Fig. 1*L*). Thus, basal NFE2L1 protein that escapes proteasomal degradation appeared sufficient to inhibit ferroptosis, with greater accumulation of this transcription factor not conferring any additional protection.

### Protein Sequence-Edited NFE2L1 Promotes Ferroptosis Resistance

NFE2L1 is subject to complex post-translational regulation, including N-glycosylation and subsequent deglycosylation by NGLY1 (26) (Fig. 2*A*). NGLY1-mediated deglycosylation is of special interest as this enzyme is mutated in an ultrarare developmental disorder of unknown etiology called Ngly1 deficiency. Accordingly, we examined whether loss of NGLY1, and subsequent NFE2L1 misprocessing, altered NFE2L1 function and ferroptosis sensitivity. We generated clonal *NGLY1* gene disrupted cell lines expressing nuclear mKate2 (i.e. NGLY1^LOF1/2,N^). We observed no detectable NGLY1 protein, reduced NFE2L1 protein abundance, and a higher molecular weight NFE2L1 protein species corresponding to its glycosylated form in both NGLY1^LOF1/2^ cell lines (Fig. 2*B*). However, since we could not confirm the disruption of the *NGLY1* genomic locus via DNA sequencing we refer to these clones as loss of function (LOF) rather than KO. Because there remained some deglycosylated NFE2L1 in the NGLY1^LOF1/2^ (Fig. 2*B*) lines we infer that the NGLY1^LOF1/2,N^ cell lines may be NGLY1 hypomorphs rather than true knockouts. Nevertheless, both NGLY1^LOF1/2,N^ cells showed enhanced sensitivity to erastin2-induced cell death, which was confirmed to be exclusively ferroptotic using Fer-1 co-treatment (Fig. 2*C*).

**Figure 2.**
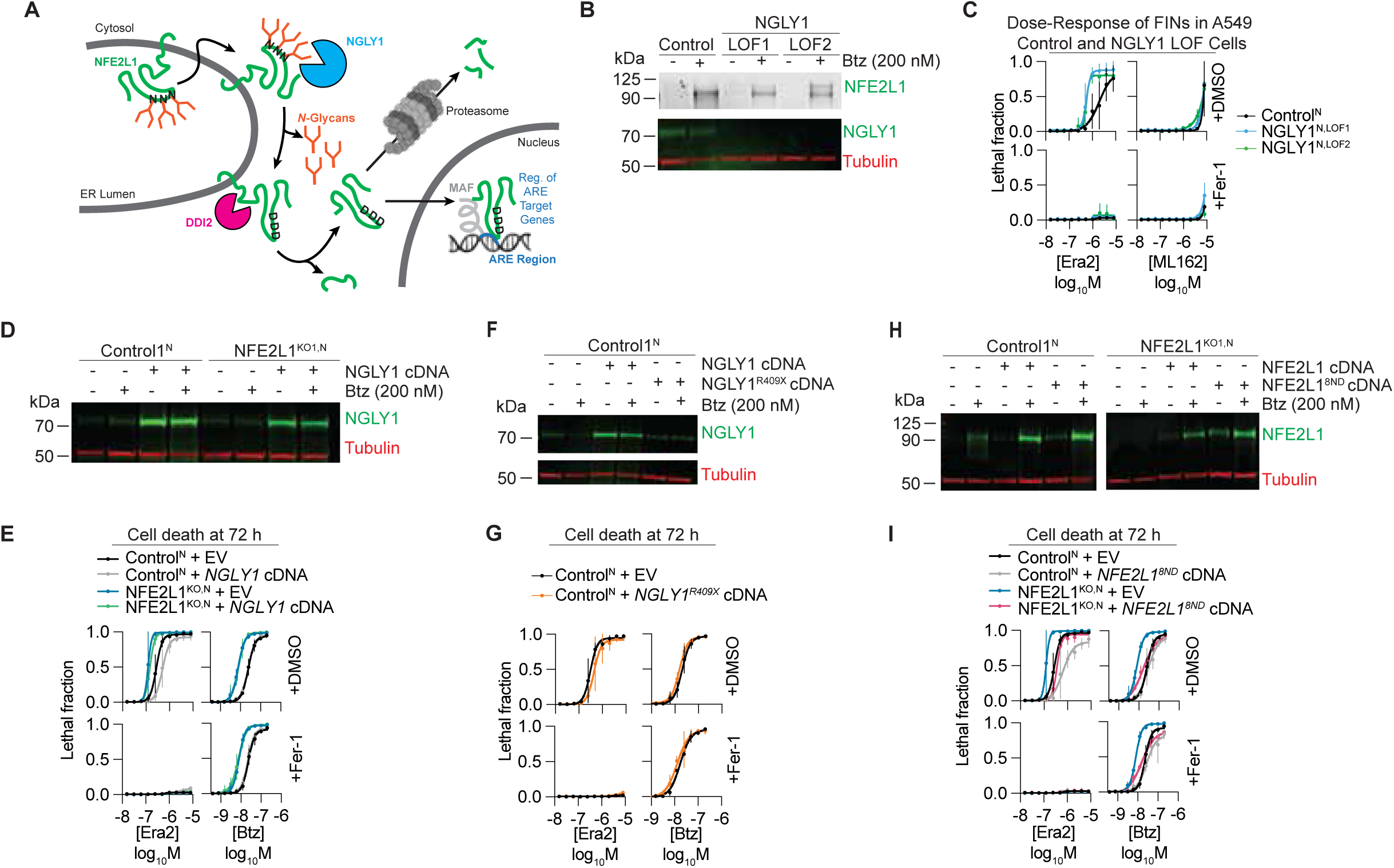
NGLY1-mediated protein sequence editing of NFE2L1 regulates ferroptosis. (*A*) Schematic of NFE2L1 processing and activation. (*B*) NFE2L1 and NGLY1 protein levels in Control^N^ and NGLY1^KO1/2,N^ cell lines. (*C*) Ferroptosis sensitivity in Control^N^ and NGLY1^KO1/2,N^ cell lines treated with a 10-point, 2-fold dose-response of erastin2 and ML162. (*D*) NGLY1 protein levels in Control1^N^ and NFE2L1^KO1,N^ cells transduced with EV or NGLY1 cDNA. (*E*) Ferroptosis sensitivity in Control1^N^ and NFE2L1^KO1,N^ cells transduced with EV or NGLY1 cDNA. (*F*) NGLY1 protein levels in Control1^N^ cells transduced with EV, NGLY1, or NGLY1^A1201T^ cDNA. (*G*) Ferroptosis sensitivity in Control1^N^ cells transduced with EV, NGLY1, or NGLY1^A1201T^ cDNA. (*H*) NFE2L1 protein levels in Control1^N^ or NFE2L1^KO1,N^ cells transduced with EV, NFE2L1, or sequence edited NFE2L1^8ND^ cDNA. (*I*) Ferroptosis sensitivity in Control1^N^ and NFE2L1^KO1,N^ cells transduced with EV, NFE2L1, or sequence edited NFE2L1^8ND^ cDNA. Data in (*C*) represent mean ± S.D. of three biological replicates. Data in (*E,I*) represent mean ± S.D. of four biological replicates.

NGLY1 and NFE2L1 operate within the same pathway to regulate bortezomib-induced cell death (26). We tested whether NGLY1 and NFE2L1 likewise operated in the same pathway to regulate ferroptosis. First, we generated Control1^N^ and NFE2L1^KO1,N^ cell lines transduced with lentivirus harboring *NGLY1* cDNA (Fig. 2*D*). Control1^N^ cells overexpressing NGLY1 were better protected against erastin2-induced ferroptosis compared to Control1^N^ cells expressing an empty vector (EV) control (Fig. 2*E*). By contrast, sensitivity to erastin2 in NFE2L1^KO1,N^ cells was unchanged when NGLY1 was overexpressed (Fig. 2*E*). Interestingly, we observed that NGLY1 overexpression had no effect on bortezomib-induced cell death in these cells (Fig. 2*E*). Unlike wild-type NGLY1, expression of an Ngly1 deficiency-associated NGLY1 disease variant resulting in a premature stop codon, NGLY1^R409X^ had no ability to accumulate via western blot or inhibit ferroptosis in Control1^N^ cells (Fig. 2*F,G*). These data suggested that NGLY1 operates upstream of NFE2L1 to regulate sensitivity to ferroptosis.

In *C. elegans*, NGLY1/PNG-1-mediated deglycosylation of NFE2L1/SKN-1A results in a protein sequence editing event where asparagine residues are converted to aspartic acid residues (28). This Asn→Asp editing is required for proper NFE2L1/SKN-1A function to protect against bortezomib-induced cell death (28). In *C. elegans*, SKN-1A has four potential sites of N-glycosylation. The human NFE2L1 protein is considerably larger and contains at least eight potential sites of N-glycosylation (predicted by conserved N-glycosylation motifs NxS/T, where x is any amino acid except proline). Which, if any, of these sites were important for NFE2L1 ferroptosis regulation was unclear. To investigate, we stably expressed a fully ‘edited’ NFE2L1 construct where all eight predicted glycosylated Asn residues were converted to Asp (NFE2L1^8ND^), mimicking NGLY1-dependent processing. We expressed wild-type NFE2L1 and NFE2L1^8ND^ in both Control1^N^ and NFE2L1^KO1,N^ cells. Like wild-type NFE2L1, NFE2L1^8ND^ accumulated following bortezomib treatment (Fig. 2*H*). Notably, NFE2L1^8ND^ appeared more stable than wild-type NFE2L1, as we were able to detect significant levels of the mutant protein by western blot under uninduced (e.g. DMSO-treated) conditions (Fig. 2*H*). We infer that the presence of N-glycans on NFE2L1 may promote its degradation. Compared to cells expressing wild-type NFE2L1, both Control1^N^ and NFE2L1^KO1,N^ cells expressing NFE2L1^8ND^ displayed enhanced resistance to cell death induced by bortezomib or erastin2 (Fig. 2*I*). These results suggest that deglycosylation-mediated protein sequence editing of NFE2L1 may be necessary for cell death resistance.

### The NGLY1/NFE2L1 Pathway Regulates GPX4 Levels

We next sought to determine the factors downstream of NFE2L1 that regulate ferroptosis sensitivity. We first performed unbiased RNA sequencing (RNA-seq) analysis of A549 Control1 and NFE2L1^KO1/2^ cells. Bortezomib was used as a positive control, and we confirmed that this treatment caused the induction of numerous proteasomal subunit genes in Control1 cells that were absent in NFE2L1^KO1/2^ cells (Figure S*1A*). However, given our results above suggesting that basal NFE2L1 expression was sufficient to regulate ferroptosis sensitivity, we focused on differences in gene expression between vehicle (DMSO)-treated Control1 and NFE2L1^KO1/2^ cells. In total, the expression of 281 genes were significantly (p_adj_ < 0.05) altered across two biological replicates in NFE2L1^KO1^ cells compared to Control1, with 247 genes significantly altered in NFE2L1^KO2^ cells compared to Control1. The expression of 97 genes were significantly altered (p_adj_ < 0.05) in both knockout lines, with 56 genes increased and 41 genes decreased (Fig. 3*A*). Relevant to ferroptosis, we did not observe decreased expression of *SLC7A11*, ferroptosis suppressor protein 1 (*FSP1/AIFM2*), or glutathione biosynthetic genes (*GCLC, GCLM, GSS*) in NFE2L1^KO1/2^ versus Control1 cells. However, we did observe a decrease in *GPX4* mRNA levels in both KO cell lines versus controls, which we subsequently confirmed by RT-qPCR (Fig. 3*B*).

**Figure 3.**
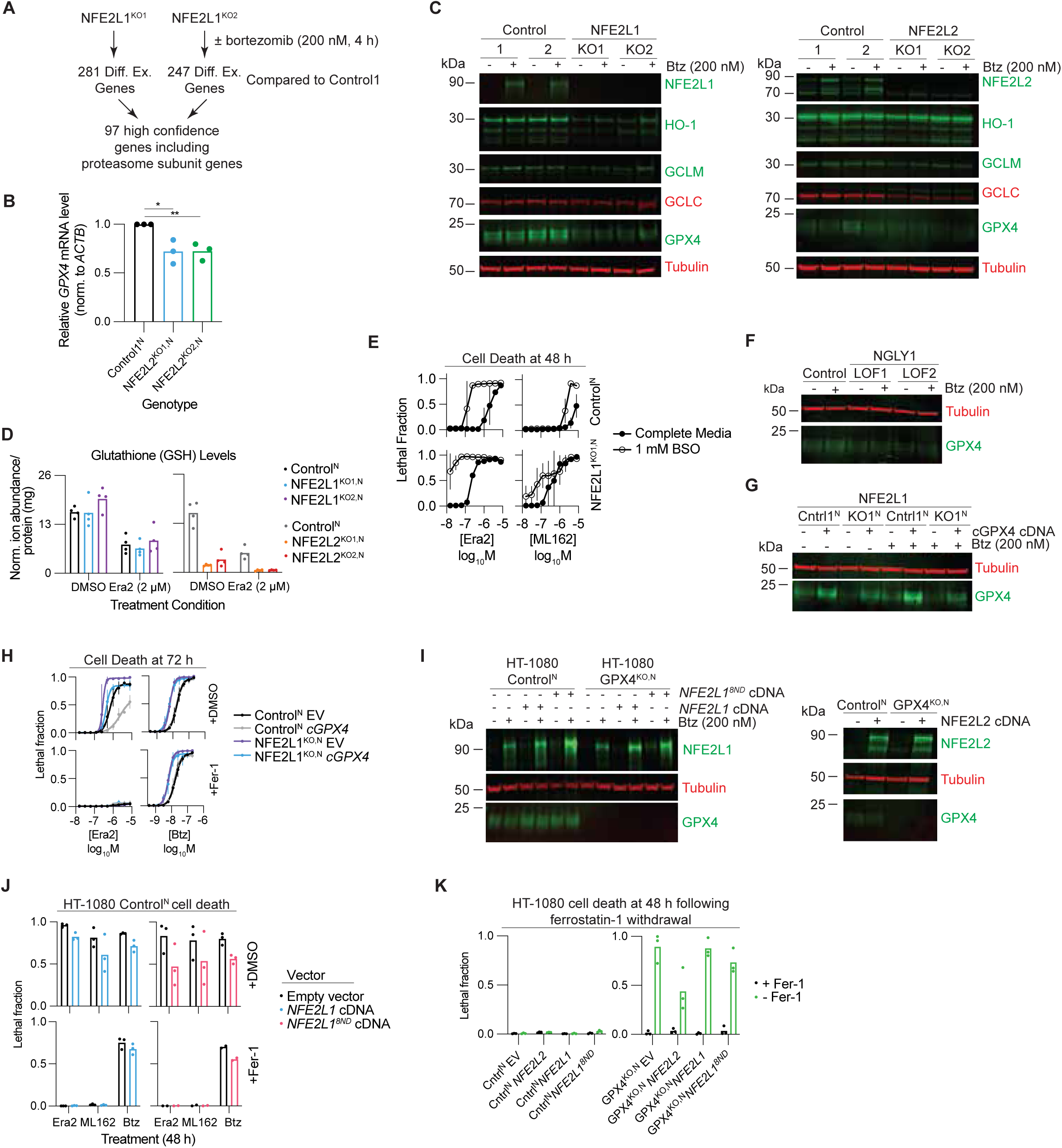
NFE2L1 and NFE2L2 regulate GPX4. (*A*) Overview of unbiased RNA-sequencing analysis. (*B*) RT-qPCR for *GPX4* mRNA levels in Control1^N^ or NFE2L1^KO1/2,N^ cells. (*C*) Western blot analysis of genes identified in RNA-seq and of genes that regulate ferroptosis in Control^N^ and NFE2L1^KO1/2,N^ or NFE2L2^KO1/2,N^ cell lines. (*D*) Glutathione levels in Control^N^ and NFE2L1^KO1/2,N^ or NFE2L2^KO1/2,N^ cell lines treated with vehicle (DMSO) or erastin2 (4 µM) for 4 h. (*E*) Ferroptosis sensitivity in Control1^N^ and NFE2L1^KO1,N^ cells in the presence or absence of 24 h pretreatment with BSO. (*F*) GPX4 protein levels in Control1^N^ and NGLY1^KO1/2,N^ cells. (*G*) GPX4 protein levels in Control1^N^ and NFE2L1^KO1,N^ cells transduced with EV or cGPX4 cDNA. (*H*) Ferroptosis sensitivity in Control1^N^ and NFE2L1^KO1,N^ cells transduced with EV or cGPX4 cDNA. (*I*) NFE2L1 and NFE2L2 protein levels in HT-1080 Control^N^ and GPX4^KO,N^ cell lines transduced with EV, NFE2L1, or NFE2L2 cDNA. (*J*) Effect of NFE2L1 or NFE2L2 overexpression on HT-1080 Control^N^ and GPX4^KO,N^ cells upon Fer-1 withdrawal. (*K*) Effect on NFE2L1 overexpression in HT-1080 Control^N^ cells on erastin2 (125 nM), ML162 (250 nM), or bortezomib (100 nM) induced cell death. Data in (*D*) represents four biological replicates. Data in (*E*) represents mean ± S.D. of two biological replicates. Data in (*H,J,K*) represent mean ± S.D. of three biological replicates.

To explore further the impact of NFE2L1 disruption we examined the levels of various proteins by western blotting (Fig. 3*C*). In this experiment we also compared the effects of NFE2L1 disruption to NFE2L2 disruption, to see whether these two transcription factors had any important differential effects on the expression of key proteins. For this analysis we included canonical NFE2L2 target gene products (e.g. HMOX1), key ferroptosis-associated gene products, some of which are known NFE2L2 targets (e.g. GCLC, GCLM), and other ferroptosis-associated genes (GPX4). Disruption of NFE2L1 resulted in loss of heme oxygenase HO-1 (encoded by *HMOX1*) and GCLM, with little effect on the levels of GCLC (Fig. 3*C, left*). Consistent with the decreased *GPX4* mRNA in NFE2L1^KO^ lines we observed a marked decrease in GPX4 protein (Fig. 3*C*). By contrast, disruption of NFE2L2 resulted in strong loss of GCLC, consistent with previous analyses of mouse embryonic fibroblasts (20), but surprisingly little change in HO-1, GCLM and GPX4 levels (Fig. 3*C, right*). Bortezomib treatment, included as a positive control for NFE2L1 accumulation in Control1/2 cell lines, did not consistently alter the expression of HO-1, GCLM, GCLC or GPX4 in Control or NFE2L1/2^KO^ cell lines (Fig. 3*C*). Thus, basal NFE2L1 expression positively regulates the abundance of certain antioxidant and ferroptosis-associated proteins in a manner distinct from NFE2L2.

A notable observation concerned differences between gene disrupted cell lines in the regulation of GCLC and GCLM. GCLC and GCLM form GCL, which catalyzes the first and rate-limiting step in GSH synthesis (35). GCLC is the catalytic subunit while GCLM is a regulatory subunit. Based on the observed changes in protein expression, we hypothesized that GSH levels would be more strongly reduced in cells lacking NFE2L2 than NFE2L1. Consistent with this hypothesis, we observed using liquid chromatography/ mass spectrometry (LC-MS) analysis that GSH levels were not reduced in NFE2L1^KO1/2^ cells, while loss of NFE2L2 greatly reduced intracellular GSH, almost to the levels obtained following treatment of Control cells with erastin2 (Fig. 3*D*). We hypothesized that the greater sensitization to ferroptosis we observed previously in NFE2L2^KO^ versus NFE2L1^KO^ cells (Fig. 1*D,H*) could be explained in part by these difference in basal GCLC expression and GSH abundance. Consistent with this hypothesis, pretreating NFE2L1^KO,N^ cells for 24 h with the GCLC inhibitor buthionine sulfoximine (BSO) sensitized NFE2L1^KO,N^ cell lines to erastin2-induced ferroptosis to levels similar to that observed with NFE2L2 disruption (Fig. 3*E*, Fig. 1*D*). Thus, NFE2L1 and NFE2L2 have different effects on basal glutathione metabolism, which correlates with variation in ferroptosis sensitivity.

As GSH levels were not strongly reduced in cells lacking NFE2L1, this left unanswered how these cells were sensitized to ferroptosis. Notably, we observed a striking decrease in GPX4 protein levels in NFE2L1^KO1/2^ cells compared to Control1/2 cells (Fig. 3*C*). GPX4 levels were also reduced in NGLY1^LOF1/2,N^ lines compared to Control^N^ cells (Fig. 3*F*). Accordingly, we hypothesized that loss of GPX4 protein upon NGLY1/ NFE2L1 pathway disruption sensitized cells to ferroptosis. To test this hypothesis, we generated Control1^N^ and NFE2L1^KO1,N^ lines stably overexpressing the cytosolic isoform of GPX4 (cGPX4). We observed that cGPX4 overexpression was sufficient to selectively reverted sensitivity to erastin2-induced ferroptosis, with no effect on bortezomib toxicity (Fig. 3*G,H*). Thus, cGPX4 expression is sufficient to inhibit ferroptosis in cells lacking NFE2L1.

NFE2L1 can regulate the expression of numerous antioxidant genes, any one of which could be required to suppress ferroptosis (20) (Fig. 3*C*). To test whether GPX4 alone was necessary for suppression of ferroptosis we transduced *KEAP1* wild-type HT-1080 fibrosarcoma Control^N^ and GPX4^KO,N^ cells with vectors directing the expression of NFE2L1 or, as a comparison, NFE2L2 (Fig. 3*I*). Due to strong proteasome-mediated negative regulation, cells transduced with the WT NFE2L1 cDNA expressed only slightly elevated levels of this protein compared to empty vector (EV) controls; however expression of NFE2L1^8ND^ accumulated to higher levels in both bortezomib- and vehicle-treated cells (Fig. 3*I*). Consistent with a model for NFE2L1 regulating GPX4 levels, both transduction with WT NFE2L1 and NFE2L1^8ND^ cDNA, but not NFE2L2 cDNA, increased levels of GPX4 (Fig. 3*I*). Moreover, consistent with results obtained in A549 cells (Fig. 1*H*), HT-1080 Control^N^ cells overexpressing NFE2L1 or NFE2L1^8ND^ were less sensitive to ferroptosis induced by erastin2 or ML162, as well as to non-ferroptotic cell death induced by bortezomib (Fig. 3*J*). We then examined GPX4^KO^ cells, which must be cultured continually in medium containing Fer-1 to prevent cell death. In GPX4^KO,N^ cells cultured without Fer-1, NFE2L1 overexpression no longer protected against ferroptosis (Fig. 3*K*) with only marginal protection afforded by NFE2L1^8ND^. By contrast, GPX4^KO,N^ cultured without Fer-1 were partially protected from ferroptosis by NFE2L2 overexpression (Fig. 3*K*). Thus, GPX4 is necessary and sufficient for the NGLY1/NFE2L1 pathway to inhibit ferroptosis, while NFE2L2 can inhibit ferroptosis in a manner that is partially GPX4 independent.

### NFE2L2 can compensate for disruption of the NGLY1/NFE2L1 pathway

The functional interplay between NFE2L1 and NFE2L2 is of great interest. The embryonic lethality of *Nfe2l1* knockout mice indicates that *Nfe2l2* cannot fully compensate for the absence of the former gene (16, 17). In cancer cells, we observed distinct patterns of genetic dependence which also suggest unique functions for both proteins. Whether NFE2L2 can compensate for loss of NFE2L1 is especially relevant in the context of NGLY1 disruption, as occurs in NGLY1 deficiency. Accordingly, we first examined NFE2L2 levels in A549 NFE2L1^KO1^ cells, and vice versa, to search for evidence of cross-regulation. However, we observed little change in NFE2L1 protein levels in NFE2L2^KO^ cells, and only a slight decrease in NFE2L2 levels in NFE2L1^KO^ cells (Fig. 4*A*). Of note, bortezomib could increase NFE2L2 levels in both Control and NFE2L1^KO^ cell lines, suggesting that this effect did not require NFE2L1 (Fig. 4*A*).

**Figure 4.**
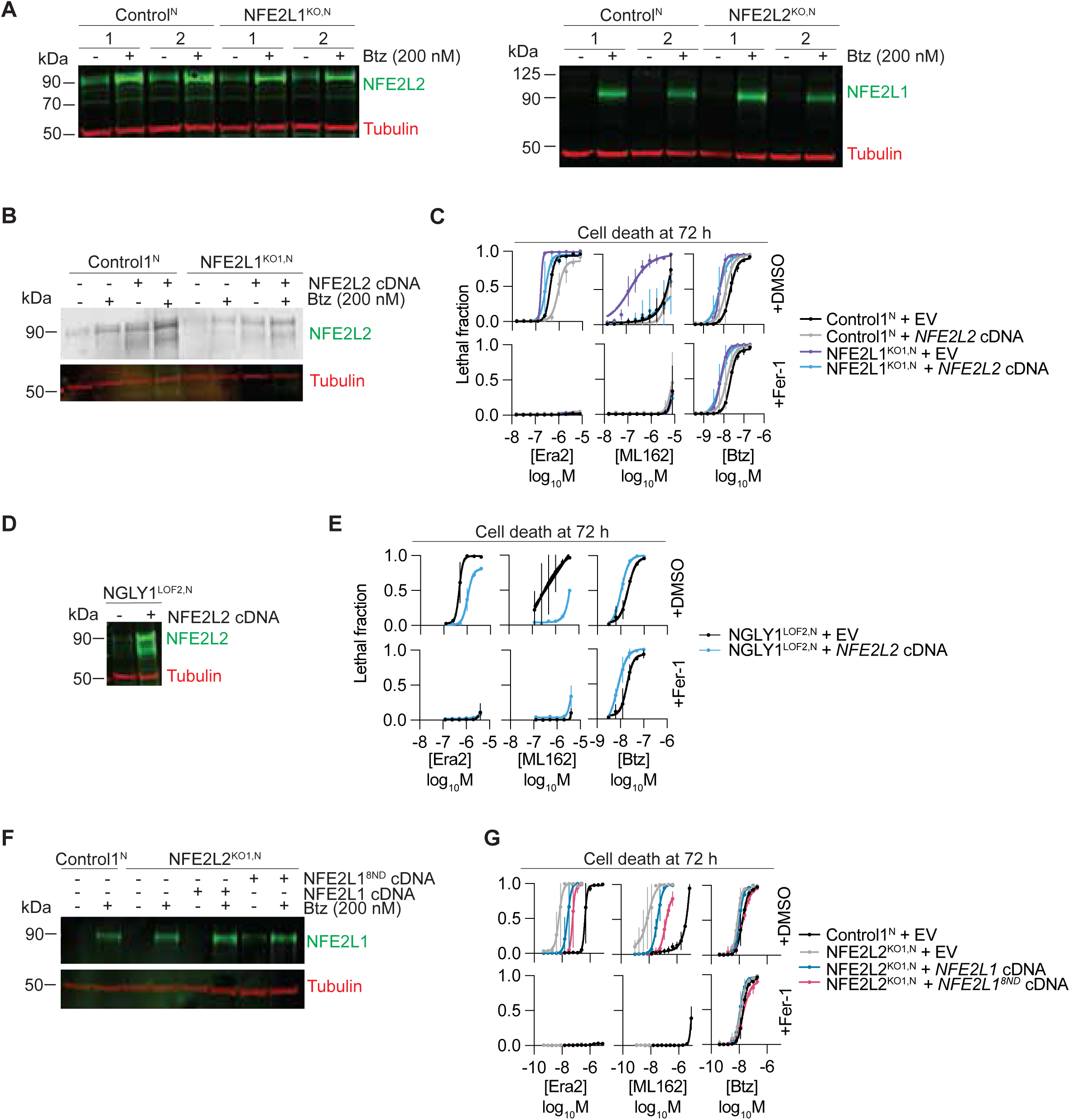
NFE2L2 and NFE2L1^8ND^ compensation to inhibit ferroptosis. (*A*) Left panel, NFE2L2 levels in Control^N^ and NFE2L1^KO,N^ cells. Right panel, NFE2L1 levels in Control^N^ and NFE2L2^KO,N^ cells. (*B*) NFE2L2 protein levels in Control1^N^ and NFE2L1^KO1,N^ cells transduced with EV or NFE2L2 cDNA. (*C*) Ferroptosis sensitivity in Control1^N^ and NFE2L1^KO1,N^ cells transduced with EV or NFE2L2 cDNA. (*D*) NFE2L2 protein levels in Control1^N^ and NGLY1^LOF2,N^ cells transduced with EV or NFE2L2 cDNA. (*E*) Ferroptosis sensitivity in Control1^N^ and NGLY1^LOF1,N^ cells transduced with EV or NFE2L2 cDNA (*F*) NFE2L1 protein levels in Control1^N^ and NFE2L2^KO1,N^ cells transduced with EV, NFE2L1, or NFE2L1^8ND^ cDNA. (*G*) Ferroptosis sensitivity in Control1^N^ and NFE2L2^KO1,N^ cells transduced with EV, NFE2L1, or NFE2L1^8ND^ cDNA. Data in (*C*) and (*G*) represent mean ± S.D. of three biological replicates. Data in (*E*) represents mean ± S.D. of two biological replicates.

Our results above suggested that NFE2L1 and NFE2L2 could regulate ferroptosis through distinct molecular mechanisms. However, it was unclear from these experiments whether NFE2L2 could functionally compensate for loss of NGLY1/NFE2L1 pathway activity, with respect to ferroptosis sensitivity. Accordingly, we generated Control1^N^ and NFE2L1^KO1,N^ cells overexpressing NFE2L2 and observed that this was sufficient to inhibit cell death caused by Era2 or ML162, but not bortezomib (Fig. 4*B,C*). NFE2L2 overexpression also was sufficient to inhibit ferroptosis in NGLY1^LOF2,N^ cells (Fig. 4*D,E*). Finally, we explored whether NFE2L1 or NFE2L1^8ND^ expression could reciprocally compensate for loss of NFE2L2. We observed that expression of wild-type NFE2L1 or NFE2L1^8ND^ in NFE2L2^KO1,N^ cells was sufficient to inhibit ferroptosis, with NFE2L1^8ND^ exhibiting a stronger effect (Fig. 4*F,G*). Thus, NFE2L2 can compensate for disruption of the NGLY1/NFE2L1 pathway, and NFE2L1 can compensate for loss of NFE2L2 function in the prevention of ferroptosis.

## Discussion

Ferroptosis sensitivity is governed by several transcription factors, including p53 (36-38), YAP1/TAZ (39, 40), and NFE2L2 (8-11). Here we show that the NGLY1/NFE2L1 pathway can also antagonize ferroptosis, in parallel to the related KEAP1/NFE2L2 pathway. Two lines of evidence suggest that NGLY1 protects cells from ferroptosis via deglycosylation of NFE2L1. First, overexpression of wild-type NGLY1 renders cells resistant to ferroptosis, but only when NFE2L1 is present. Second, the protein sequence edited mimic of NFE2L1, NFE2L1^8ND^, is sufficient to rescue ferroptosis. These results confirm and extend results obtained in *C. elegans*, suggesting that NFE2L1 protein sequence editing is crucial for the function of this transcription factor (28). In Control cells, NFE2L1^8ND^ accumulated to higher levels and more potently inhibited ferroptosis than the wild-type protein, suggesting that NGLY1-mediated deglycosylation may normally stabilize NFE2L1 protein. It was recently suggested that a ninth Asn site (Asn574) is also edited by NGLY1, and that this is essential for the function of NFE2L1 in the proteasome bounce-back response (41). We find that modification of the eight sites assayed here is sufficient to inhibit ferroptosis, and it will be interesting in future studies to elucidate whether site-specific modifications, or cellular context, dictate unique biological outcomes associated with NFE2L1 glycosylation.

We find that the NGLY1/NFE2L1 pathway basally inhibits ferroptosis by promoting the expression of GPX4: disruption of *NGLY1* or *NFE2L1* reduces GPX4 protein expression and overexpression of GPX4 is sufficient to rescue ferroptosis sensitivity in *NFE2L1* mutants. NFE2L1 regulates transcription by dimerizing with small Maf cofactors to occupy antioxidant response elements in the promoter region of target genes (42). How NFE2L1 and related family members selectively modulate specific ARE-containing genes is not well understood (43). We note that the genomic *GPX4* locus contains two candidate ARE-like sequences and an NF-E2/AP1 consensus motif in its promoter region which may mediate NFE2L1 binding. Proteasome inhibition can enable NFE2L1 to accumulate to high levels and drive a proteasome bounce back response (22). Interestingly, however, NFE2L1 accumulation caused by proteasome inhibition did not boost protection against ferroptosis. Thus, the basal pool of stabilized NFE2L1 that escapes proteasomal degradation is sufficient maintain GPX4 expression and inhibit ferroptosis constitutively. It is interesting to consider that the NGLY1/NFE2L1 pathway emerges from our analysis as an important regulator of sensitivity to both apoptotic and non-apoptotic lethal stimuli.

In humans, *NGLY1* mutation causes a devastating multi-organ disorder that is presently uncurable. Whether the symptoms associated with NGLY1 deficiency can be explained by the death of individual cells is presently unclear. At the cellular level, NGLY1 disruption has been associated with defects in mitochondrial homeostasis and inflammatory responses (44, 45), AMPK signaling (46), plasma membrane transporter function (47, 48) and BMP4 signaling (49). However, analysis of an *Ngly1* deficient *D. melanogaster* model suggest a role for enhanced oxidative stress in this disease (50). Among other observations, we find that the disease associated NGLY1^R409X^ mutant is unable to inhibit ferroptosis. It is important to note that our studies were performed largely in transformed cells. Nonetheless, our results are consistent with the possibility that enhanced oxidative stress contributes to the etiology of NGLY1 deficiency by promoting ferroptosis. Our results support the notion that compounds which cause the accumulation of NFE2L2 could provide a therapeutic benefit under conditions of NGLY1/NFE2L1 pathway dysfunction (51). Indeed, we speculate that the beneficial effects of NFE2L2 activating compounds observed previously in worm and fly models of NGLY1 deficiency (51) may be due in part to inhibition of ferroptosis.

## Materials and Methods

### Chemicals and reagents

A 261-member bioactive compound library were obtained from Selleck Chemicals (Cat# L2000), formatted as described previously (6), and stored at -80°C. Erastin2 and ML162 were synthesized by Acme Bioscience. Chemical used (supplier) were: DMSO, ferrostatin-1 (Sigma-Aldrich Corporation); Q-VD-OPh, bortezomib, and buthionine sulfoximine (BSO, Thermo-Fisher Scientific); camptothecin, 1*S*,3*R*-RSL3 (RSL3), and bazedoxifene (Selleck Chemicals). BSO was dissolved directly into cell media. All other drugs were prepared as stock solutions in DMSO. Stock solutions were stored at -20°C.

### Cell Culture

A549 (CCL-185, gender: male) and HT-1080 (CCL-121, gender: male) cells were obtained from ATCC, expanded for one passage then frozen down in multiple aliquots used for all subsequent experiments. Mouse embryonic fibroblasts (gender: not specified) were derived from knockout animals and immortalized using SV40, and were a kind gift from Tadashi Suzuki (RIKEN, Japan). Paired control and NGLY1^KO^ HEK293 and HepG2 cells were a kind gift from Grace Science, LLC (Menlo Park, CA). A549 (gender: male) CRISPR/Cas9 Control1/2 cells and NFE2L1^KO1/2^, NGLY1^LOF1/2^, NFE2L2^KO1/2^, and HT-1080 GPX4^KO1/2^ cells expressing nuclear-localized mKate2 (denoted by an additional superscript ‘N’) were generated as described below. Polyclonal populations of these cell lines were generated from the respective parental cells via transduction with the NucLight Red lentivirus, which directs the expression of nuclear-localized mKate2 (Essen BioSciences/Sartorius). Polyclonal mKate2 expressing cells were selected for using FACS (Stanford Shared FACS Facility). All cells were grown in Dulbecco’s modified Eagle medium (DMEM, Cat# MT-10-013-CV, Thermo Fisher Scientific, Waltham, MA) supplemented with 10% fetal bovine serum (FBS, Cat# 26140-079, Gibco), and 0.5 U/mL Pen/Strep (P/S, Cat# 15070-063, Gibco). HT-1080 cells were additionally supplemented with 1% MEM nonessential amino acids (Cat# 11140-050, Life Technologies). 1X PBS (Cat# 97062-338) is from VWR and trypsin (Cat# 25200114) are from Gibco. For all experiments, cells were trypsinized and counted using a Cellometer Auto T4 cell counter (Nexcelom, Lawrence, MA, USA).

### Genotype drug sensitivity analysis

Data for cell line *KEAP1* mutation status and sensitivity to erastin and RSL3 (from the CTD^2^ program) were downloaded from the DepMap portal (https://depmap.org/portal/) July 19^th^, 2021.

### Generation of clonal CRISPR/Cas9 KO Cell

The following sgRNA primer sets were used to generate clonal *NFE2L1, NGLY1, NFE2L2*, and *GPX4* gene disrupted cell lines, which we refer to as either knockouts (KO) or loss of function (LOF): *NFE2L1* (exon 2), forward: 5’-CACCgTTTCTCGCACCCCGTTGTCT-3’ and reverse: 5’-AAACAGACAACGGGGTGCGAGAAAc-3’; *NGLY1* (exon 2), forward: 5’-CACCgAAATATAGATCCATCCGGAT-3’ and reverse: 5’-AAACATCCGGATGGATCTATATTTc-3’; *NFE2L2* (exon 2), forward: 5’-CACCgATTTGATTGACATACTTTGG-3’ and reverse: 5’-AAACCCAAAGTATGTCAATCAAATc-3’; *GPX4* (exon 3), forward: 5’-CACGCCCGATACGCTGAGTG-3’ and reverse: 5’CACTCAGCGTATCGGGCGTGc-3’. SgRNAs were cloned into pSpCas9(BB)-2A-GFP (a kind gift from Dr. Jan Carette (Stanford University School of Medicine), as described (52). Briefly, primers were annealed by heating to 95°C, then cooled by 2.5°C/minute to 25°C. The resulting oligo duplex was diluted 1:200 in ddH_2_O and used for cloning into the pSpCas9(BB) plasmid. The ligation reaction was as follows: 100 ng pSpCas9(BB), 2 μL of the diluted oligo duplex, 2 μL of 10x FastDigest buffer (Thermo Scientific), 1 μL 10 mM DTT, 1 μL 10 mM ATP, 1 μL FastDigest BpiI (Cat# D1014, Thermo Scientific), 0.5 μL T4 DNA ligase (Cat# EL0014, Thermo Scientific), and 12.5 µL of ddH_2_O to produce a final reaction volume of 20 µL. The ligation reaction was incubated for 1 h [(37°C, 5 min; 21°C, 5 min) x 6 cycles]. 2 μL of the product was used to transform competent DH5α cells on LB-amp (100 µg/mL) plates. Plasmid DNA was extracted using a QIAGEN spin column (Cat# 27106, QIAGEN, Hilden, Germany) and confirmed via Sanger sequencing. Cells were transfected with 2.5 µg DNA using Opti-MEM I Reduced Serum Medium (Cat# 31985-062, Life Technologies) and Lipofectamine LTX Reagent with PLUS Reagent (Cat# 15338-100, Life Technologies), according to the manufacturer’s instructions. The next day, medium was replaced, and the cells allowed to recover for 24 h. Transfected cells were then trypsinized, washed in 1X PBS, and sorted for GFP+ cells with a Carmen Cell Sorter (BD Biosciences) (Stanford Shared FACS Facility) into a 96-well plate containing either DMEM containing 30% FBS and 0.5 U/mL P/S. Cells were incubated at 37°C until distinct clones appeared. Individual clones were amplified in normal (10% FBS) medium to produce enough cells for protein analysis. Clones were validated by western blot to confirm loss of protein expression. HT-1080 GPX4^KO1/2^ clones were cultured in medium supplemented in 1 µM Fer-1 to inhibit ferroptosis.

### Generation of stable overexpression cell lines

The cDNA of human NGLY1 was subcloned from a pLex-NGLY1 construct, described previously (26), into the lentiviral expression plasmid, pLenti-CMV-Puro DEST (Addgene plasmid #17452, from Eric Campeau and Paul Kaufman) via Gibson assembly using a commercial kit (Cat# E2611S, NEB). Lentiviral constructs for NFE2L1 and protein sequence edited NFE2L1^8ND^ were synthesized (Twist Biosciences, South San Francisco, CA). The open reading frame for NFE2L2 was obtained from Dr. Aaron Gitler as a pDONR plasmid from the human ORFeome library. The NFE2L2 ORF was cloned into the pLenti-CMV-Puro vector using a Gateway LR Clonase II kit (Cat# 11791-020, Thermo Fisher Scientfic, Waltham, MA). The cytosolic GPX4 (cGPX4) construct pLenti-CMV-Hygro-cGPX4 was a kind gift from James Olzmann. pLenti-CMV-Puro was used as an empty vector control for comparison in overexpression experiments. All plasmids were verified by Sanger sequencing. Lentiviruses were generated in HEK 293T cells using a 3rd generation lentivirus packaging system. Briefly, cells were transfected with 1 μg pLentiCMV DEST plasmid + 0.25 mg of each of three 3rd generation lentiviral packaging plasmids (pMDLg/pRRE, pRSV-Rev and pMD2.G, Addgene plasmids #12251, #12253 and #12259, respectively, from Didier Trono) using PolyJet DNA Transfection Reagent (SignaGen Laboratories, Cat# SL100688 Rockville, MD) as per the manufacturer’s instructions. Lentivirus was harvested twice (three and four days post-transfection), filtered through a 0.45 μM PVDF filter (Cat# SLHV033RS, Merck Millipore, Burlington, MA) and stored at −80°C until use. For infections, 0.5 mL of freshly thawed virus soup was mixed with polybrene (Cat# H9268, Sigma-Aldrich) to a final concentration of 8 μg/mL and used to infect A549^N^ and HT-1080^N^ cell lines. Stably transduced cell populations were selected with 1.25 μg/mL puromycin (Cat# A11138-03, Life Technologies) for two days.

### NGLY1 A1201T (NGLY1^R409X^) Mutagenesi

The NGLY1^A1201T^ mutant was generated using the Q5 Site-Directed Mutagenesis Kit (Cat# E0552S, New England Biolabs) using the pLenti-CMV-Puro-NGLY1 construct, described above, with the following primers following the manufacturer’s instructions: forward: 5’-GGTGATTGCCTGAAGAACTAAGG-3’ and reverse: 5’-TCTTCATGTTTGCAGGAATATC-3’. The resultant pLenti-CMV-Puro-NGLY1^A1201T^ construct was verified by Sanger sequencing. Lentivirus harboring this construct was generated as described above.

### Western blotting

300,000 cells/well were seeded into duplicate wells of 6-well plates (Thermo Fisher Scientific, Cat# 07-200-83) in 2 mL medium. For western blots with bortezomib induction, the day after seeding, media was replaced with 2 mL media containing either DMSO or bortezomib (200 nM final). For western blots probing NFE2L1 induction with ferroptosis inducing compounds, media was replaced with fresh media containing DMSO, erastin2 (4 µM), ML162 (4 µM), bortezomib (200 nM), erastin2 and bortezomib, or ML162 and bortezomib. After 4 h, cells were washed once with 1X PBS (VWR, Cat# 97062-338), lifted with 300 µL trypsin/well (Gibco, Cat# 25200114), and quenched with 600 µL complete medium. Duplicate wells were combined into a 2 mL collection tube (Thermo Fisher Scientific, Cat# 05-408-138) and pelleted by spinning cells at 500 g for 5 minutes. Pellets were washed with 1 mL of 1X PBS and lysed with 100 µL of 9 M urea (Cat# U5378, Sigma Aldrich). Lysates were subsequently sonicated and spun for 15 min in a centrifuge at max speed. Supernatants were collected and transferred to a fresh 1.5 mL Eppendorf tube, and protein abundance was quantified with the BCA assay. 30 µg of protein from each lysate was loaded onto a Bolt 4-12% Bis-Tris Plus SDS gel (Life Technologies) for separation for 75 min at 100 V. The gel was transferred to a nitrocellulose membrane using the iBlot2 system (Life Technologies). Membranes were probed with primary antibody overnight in a hybridization bag at 4°C. Antibody dilutions (and suppliers) are as follows: alpha-tubulin DM1A 1:5000 (Thermo Fisher Scientific, Cat# MS581P1); TCF11/NRF1/NFE2L1 D5B10 1:1000 (Cell Signaling Technology, 8052S); NGLY1 1:500 (Sigma-Aldrich, HPA036825); NRF2/NFE2L2 1:1000 (Abcam, ab137550); H5 γ-GCSc/GCLC 1:1000 (Santa Cruz Biotechnology, 390811); GCLM 1:1000 (GeneTex, GTX114075); HO-1 EPR18161-128 1:1000 (Abcam, ab189491); CPS1 EPR7493-3 1:1000 (Abcam, ab129076); GPX4 EPNCIR144 1:1000 (Abcam, ab125066). Donkey anti-rabbit secondary antibody (IRDye 800LT, LI-COR Biosciences) and donkey anti-mouse antibody (IRDye 680LT, LI-COR Biosciences) were used at 1:15000 dilution to visualize bands. Membranes were imaged using the LI-COR CLx Imaging System.

### Cell death experiments

The day prior to the start of an experiment, 1500 cells/well were seeded into a 384-well plate (Corning, Cat# 3764) in 45 µL media. The next day, the media was removed and replaced with 45 µL medium containing SYTOX Green (20 nM final) and 5 µL of 10X compound prepared in a 384-well storage plate (Thermo Fisher Scientific, Cat# AB-0781). Experiments with paired HEK293 and HepG2 control and NGLY1 KO cells were performed by seeding 7500 cells/well into a 96-well plate (Thermo Fisher Scientific, Cat# 07-200-588) in 200 µL medium. 96-well plate experiments with A549 or HT-1080 cells were seeded at a density of 4000 cells/well. For 96-well plate experiments, the day after seeding 20 µL of 10X compound was added to 180 µL of fresh media containing SYTOX Green (20 nM final). The next day the medium was removed and replaced with 180 µL medium containing SYTOX Green (20 nM final) and 20 µL of 10X compound.

### Analysis of cell death

In several small-scale experiments we examined cell death using an adapted STACK approach as described previously (34), using an IncuCyte Zoom dual color live content imaging system (Model 4459, Essen BioScience/Sartorius) housed within a Thermo tissue culture incubator (37°C, 5% CO_2_). Images were acquired using a 10x objective lens in phase contrast, green fluorescence (ex: 460 ± 20, em: 524 ± 20, acquisition time: 400 ms) and red fluorescence (ex: 585 ± 20, em: 665 ± 40, acquisition time: 800 ms) channels. For each well, images (1392 × 1040 pixels at 1.22 μm/pixel) were acquired every 2 h or 4 h for a variable length of time, depending on the experiment. Automated object detection was performed in parallel to data acquisition using the Zoom software package (V2016A/B) using a routine with the following settings (in parentheses) to count mKate2^+^ objects (Parameter adaptive, threshold adjustment: 1; Edge split on; Edge sensitivity 80; Filter area min 100 μm^2^; Mean Intensity min 3.0), SG^+^ objects (Parameter adaptive, threshold adjustment: 1; Edge split on; Edge sensitivity -5; Filter area min 20 μm^2^, maximum 750 μm^2^; Eccentricity max 1.0; Mean Intensity min 1.0), and Overlap (e.g. mKate2^+^ and SG^+^) objects (Filter area min 20 μm^2^, maximum 5000 μm^2^). Double mKate2/SYTOX Green positive counts were subtracted from live cell counts prior to the calculation of lethal fraction scores. Some experiments employed unmarked cells and SG^+^ cells only were counted as a metric of cell death.

### Small molecule library scree

The day before compound addition, 1,500 A549 Control1^N^, NFE2L1^KO1,N^, and NFE2L1^KO2,N^ cells/well were seeded into three 384-well plates in 45 μL medium. The next day, the medium was removed and replaced with medium containing SYTOX Green (20 nM final) and compounds from a freshly thawed library master stock plate (1 compound/well) were added to a final concentration of 5 μM. Plates were imaged immediately and every 4 h thereafter for a total of 96 h using the Essen IncuCyte Zoom. Counts of SYTOX Green and mKate2 objects per mm^2^ were obtained and the lethal fraction calculated as described below. The area under the curve (AUC) of lethal fraction scores across the full 96 h were calculated using the trapezoid rule in Microsoft Excel (Version 16.44). Area under the curve was normalized (e.g. nAUC) as described (53).

### RNA-sequencing

A549 Control1^N^, NFE2L1^KO1,N^, and NFE2L1^KO2,N^ cells were seeded in 6-well dishes at 200,000 cells/well. Cells were treated with either DMSO or bortezomib (200 nM) for 4 h. Cells were subsequently washed with ice-cold 1X PBS and harvested with a cell lifter over ice. Cell solution was pelleted at 500 g for 5 minutes. Following centrifugation, the supernatant was removed, and cells were lysed for RNA extraction using a Qiashredder column (Cat# 79654, Qiagen) and RNeasy Plus RNA Extraction Kit (Cat#74134, Qiagen). Two biological replicates were collected per genotype and treatment condition. Purity, concentration, and integrity of RNA was assessed by NanoDrop Spectrophotometer (Thermo Fisher Scientific) and by Eukaryote Total RNA Nano chip analysis using a Bioanalyzer (Agilent) at the Stanford Protein and Nucleic Acid Facility. RNA Integrity Number for each sample met or exceeded the minimum threshold of 6.8. Samples were then shipped on dry ice to Novogene (Sacramento, CA) for library generation and 20M read PE150 sequencing on an Illumina HiSeq 4000 Platform. Data cleanup was performed by excluding reads with: adaptor contamination; >10% of indeterminate bases, or >50% bases with a quality score ≤ 5. The remaining reads (≥98% of all reads across all conditions) were aligned to the hg19 human reference genome using STAR. At least 91% of clean reads were mapped across all conditions. Pearson correlation between biological replicates was R^2^ ≥ 0.96 for all samples. Differential gene expression was calculated using DESeq 1.10.1. We selected all protein-coding genes with read counts above 0.5 FPKM in all conditions and with significant alterations (adjusted *P* value < 0.05). All RNA sequencing data was deposited to GEO with accession number GSE182461.

### RT-qPCR

300,000 Control1^N^ or NFE2L1^KO1/2,N^ cells were seeded into a 6 well plate. The next day, cells were washed twice with ice cold 1XPBS, and scraped to harvest. RNA was extracted using a Qiashredder extraction column (Qiagen, Cat# 79654) and the RNeasy Plus RNA Extraction Kit (Cat# 74134, Qiagen). cDNA was generated using the TaqMan Reverse Transcriptase Kit according to the manufacturer’s instructions (Cat# N8080234, TaqMan). Quantitative PCR reactions were prepared with SYBR Green Master Mix (Cat# 4367659, Life Technologies) and run on an Applied Biosystems QuantStudio 3 real-time PCR machine (Thermo Fisher). Relative transcript levels were calculated using the ∆∆CT method and normalized to the *ACTB* gene. The following primers were used to amplify *ACTB*: forward, 5’-CATGTACGTTGCTATCCAGGC-3’ and reverse, 5’-CTCCTTAATGTCACGCACGAT-3’; and the following primers were used to amplify GPX4: forward, 5’-AGACCGAAGTAAACTACACTCAGC-3’ and reverse, 5’-CGGCGAACTCTTTGATCTCT-3’.

### GSH measurement by LC-MS

Cells were washed with 1.5 mL/well of 0.9% saline solution in water (over ice) and then harvested with 1mL/well of 80% methanol (on dry ice) with 500 nM 13C- and 15N-labeled amino acids (Cat#MSK-A1-1.2, Cambridge Isotope Laboratories) as internal standards. Whole-cell harvest was then transferred to pre-chilled microcentrifuge tubes and stored at -80°C. On the day of analysis, samples were vortexed for 10 min at 4°C. Clear supernatant was then transferred to polypropylene autosampler vials after centrifugation for 10 min at 15,000 rpm at 4°C. Polar metabolite profiling was performed on an ID-X tribrid mass spectrometer (Thermo Fisher Scientific) with an electrospray ionization (ESI) probe. Hydrophilic interaction chromatography (HILIC) on a SeQuant® ZIC®-pHILIC 150 × 2.1 mm column (Millipore Sigma, Cat# 1504600001) coupled with a guard 20 × 2.1 mm (Millipore Sigma, Cat# 1504380001) was used to separate polar metabolites prior to mass spectrometry analysis. Mobile phase A consists of 20 mM ammonium carbonate and 0.1% ammonium hydroxide in water, and Mobile phase B is pure acetonitrile. HILIC was performed at a flow rate of 0.15 mL/min with the following gradient: linear decrease from 80-20% B from 0-20 min, linear increase from 20-80% B from 20-20.5 min, hold at 80% B from 20.5-28 min. Full scan from m/z 70-1000 with polarity switching was used for the mass spectrometer. Ion transfer tube and vaporizer temperatures were set to 275°C and 350°C, respectively. Orbitrap resolution at 120,000, RF lens at 40%, AGC target at 1×106, and maximum injection time of 80 ms were used. Positive and negative ion voltages are 3000 V and 2500 V, respectively. Sheath gas at 40 units, aux gas at 15 units, and sweep gas at 1 unit was set for the ion source. EASY-ICTM was enabled for internal mass calibration. Metabolite abundance was quantified by TracerFinder (Thermo Fisher Scientific) using an in-house library of known metabolite standards. Mass tolerance of 5 ppm was used for extracting ion chromatograms.

### HT-1080 Control^N^ and GPX4^KO^ ferrostatin-1 withdrawal experiments

8,000 HT-1080^N^ cells expressing an empty vector, NFE2L1, or NFE2L2, were seeded into four replicate wells/genotype in a 96 well plate (Thermo Fisher Scientific, Cat# 07-200-588). The seeding density of each well was 8,000 cells/well in 200 µL of media containing 1 µM Fer-1. The next day, the medium was removed, and cells were washed 3x with 200 µL of 1X PBS. After the third wash, two wells/genotype received medium containing SYTOX Green (20 nM final) ± Fer-1 (1 µM). Cell death was then determined using STACK over 48 h.

## Acknowledgments

We thank J. Carette, A. Gitler, J. Olzmann and B. Stockwell for sharing reagents. G.C.F. was supported by the Stanford ChEM-H Chemistry/Biology Interface Predoctoral Training Program. This work was also supported by the Grace Science Foundation (to C.R.B. and S.J.D), Stanford Cancer Institute Award (to M.A.-R.), and NIH (R01CA227942 and U01 CA226051-01A1 to C.R.B., R01GM122923 to S.J.D.).

---

**SI Appendix, Figure S1.**
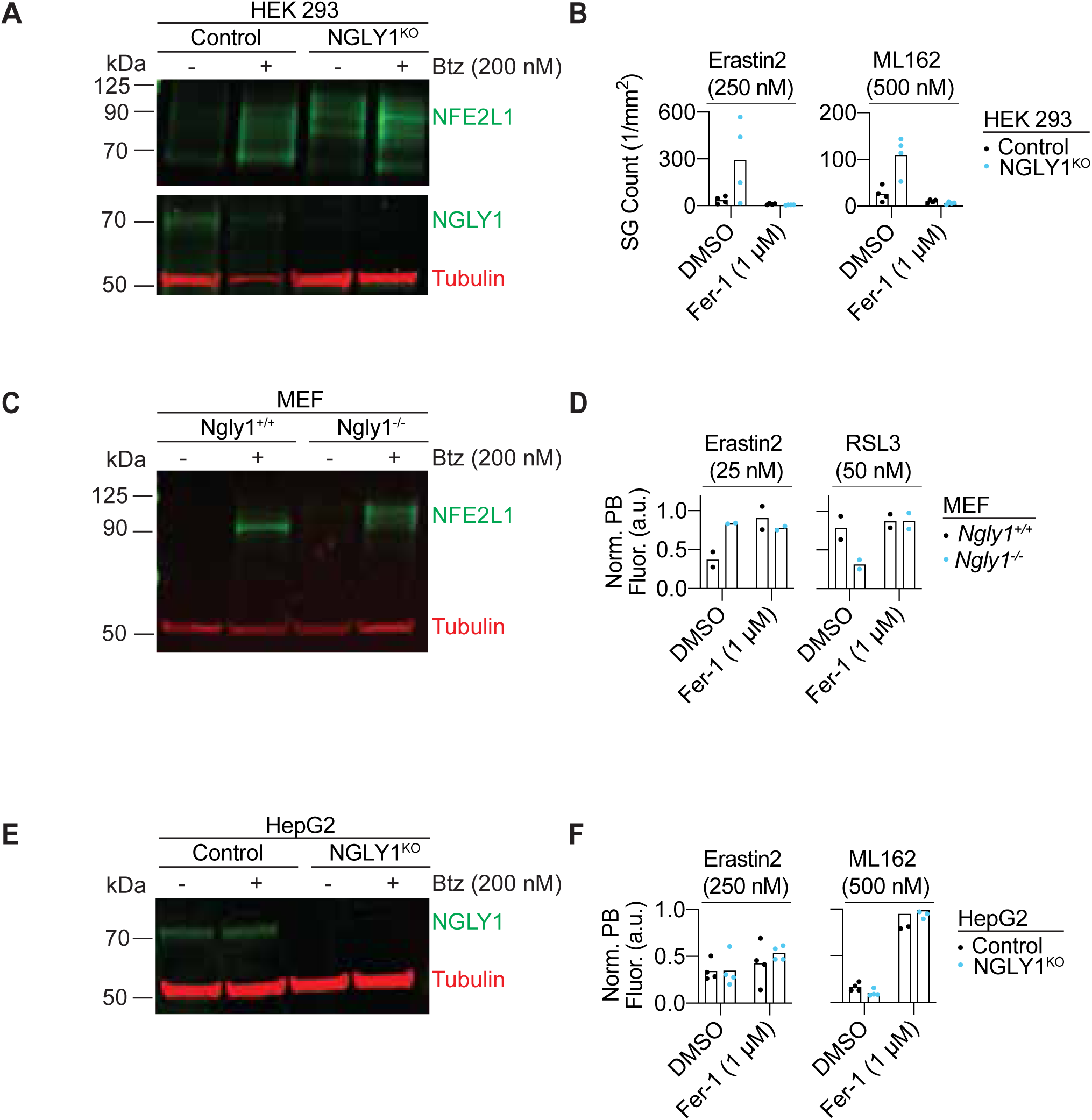
Effect of NGLY1 disruption on ferroptosis sensitivity. (*A*) Western blot of NGLY1 and NFE2L1 protein in Control and NGLY1^KO^ HEK293 cells. (*B*) Ferroptosis induction by erastin2 and ML162 in Control and NGLY1^KO^ HEK293 cells. (*C*) NFE2L1 protein in WT and NGLY1^KO^ mouse embryonic fibroblasts. (*D*) Cell viability measured with PrestoBlue for WT and NGLY1^KO^ mouse embryonic fibroblasts treated with ferroptosis inducers erastin2 and RSL3. (*E*) Western blot of NGLY1 in Control and NGLY1^KO^ HepG2 cells. (*F*) Cell viability measured with PrestoBlue for WT and NGLY1^KO^ HepG2 cells treated with ferroptosis inducers erastin2 and ML162. Data in (*B*) represent the mean ± S.D. of three biological replicates. Data in (*D*) represent the mean ± S.D. of two biological replicates. Data in (*F*) represents the mean ± S.D. of four biological replicates.

**SI Appendix, Figure S2.**
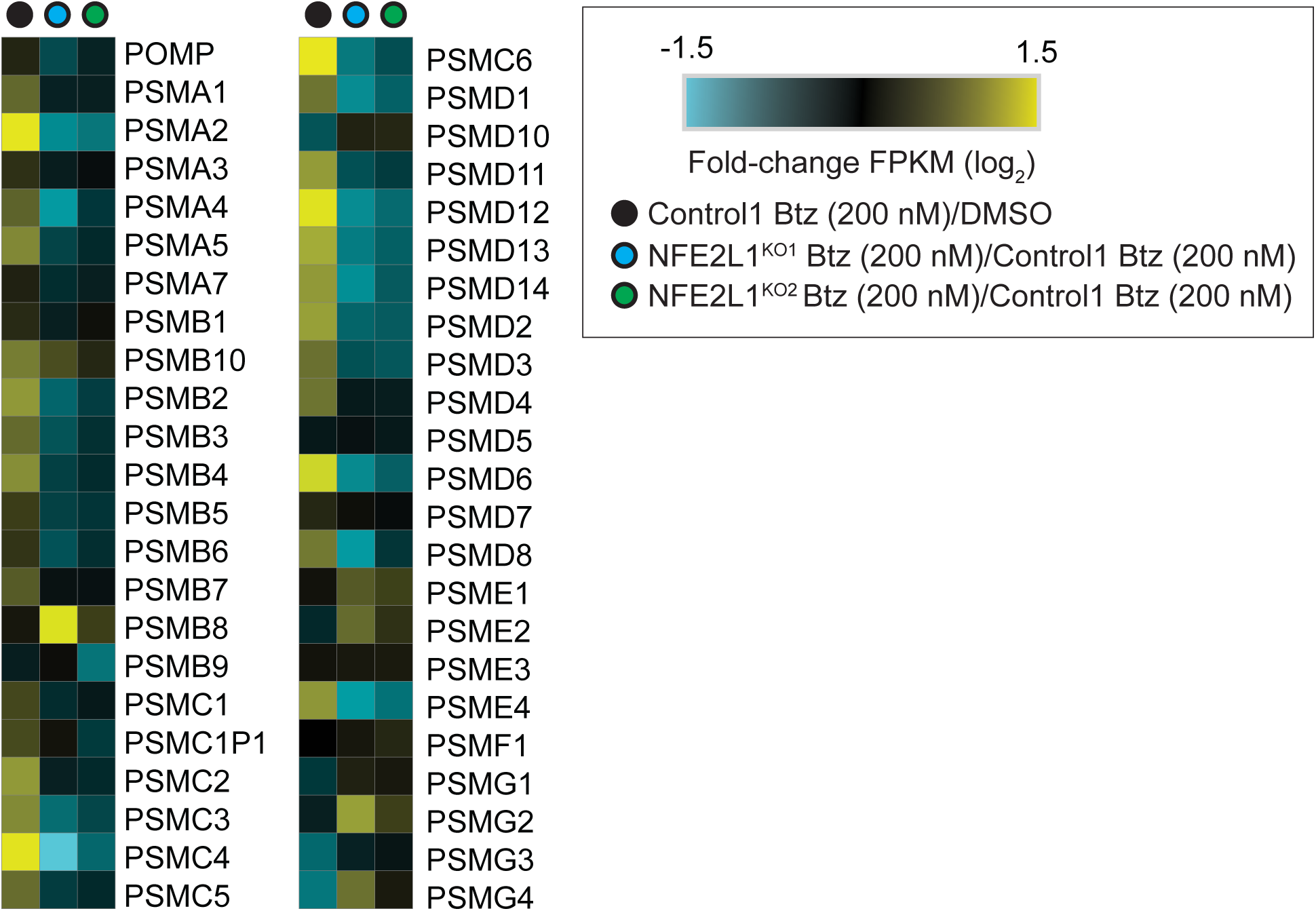
RNA sequencing analysis of proteasome genes in Control and NFE2L1^KO^ cell lines. The heatmap represents the average of FPKM fold-change of two biological replicates between Control1 and NFE2L1^KO1^ or NFE2L1^KO2^ cells. Black dot represents the gene expression of proteasome genes in Control1 cells after induction with bortezomib (200 nM) for 4 h. Blue and green dots represent the fold change between proteasome gene induction after bortezomib treatment (200 nM, 4 h) in NFE2L1^KO1^ or NFE2L1^KO2^ cells relative to Control1 cells, respectively.

**SI Appendix, Supplementary Table 1.**
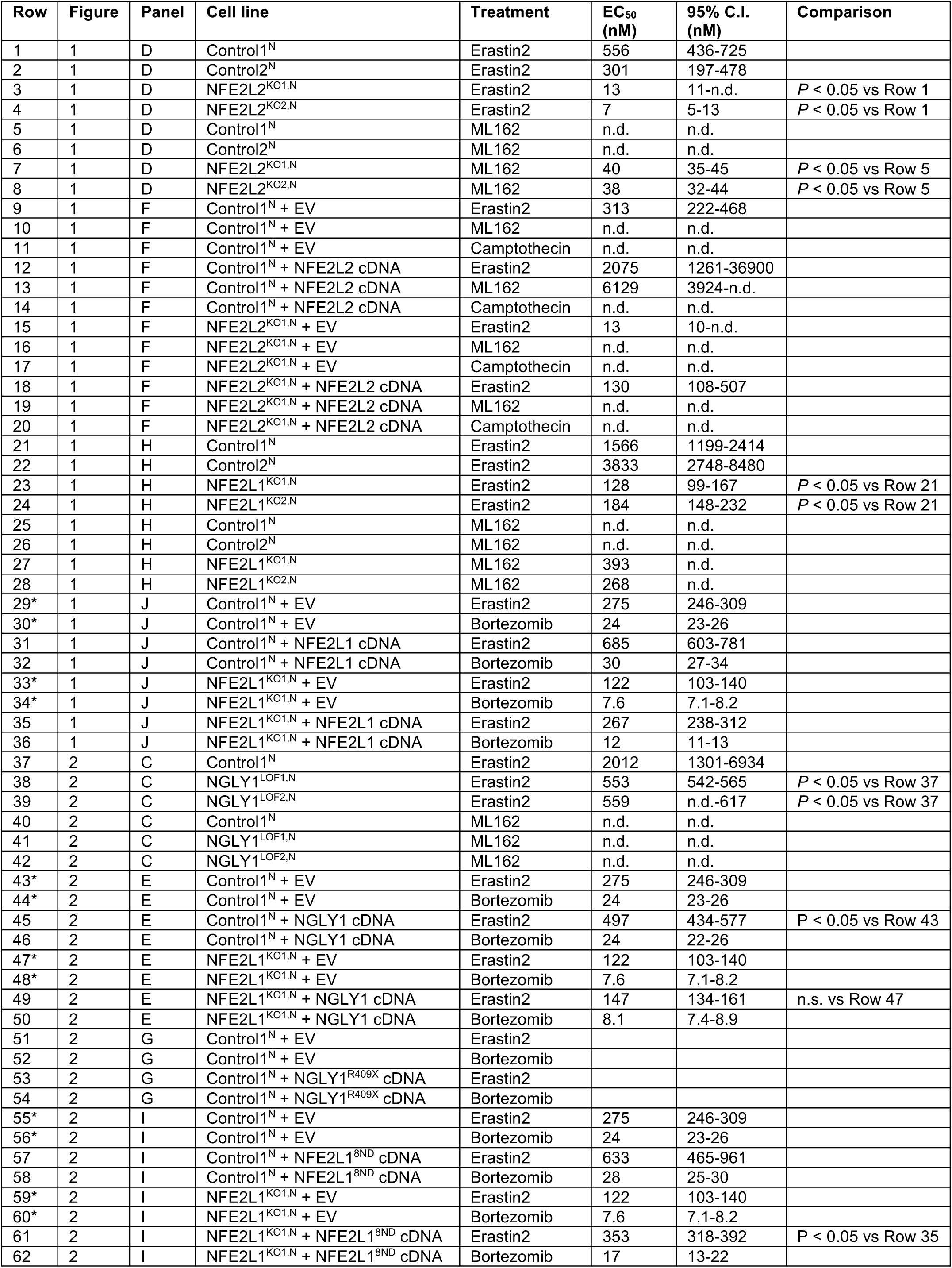
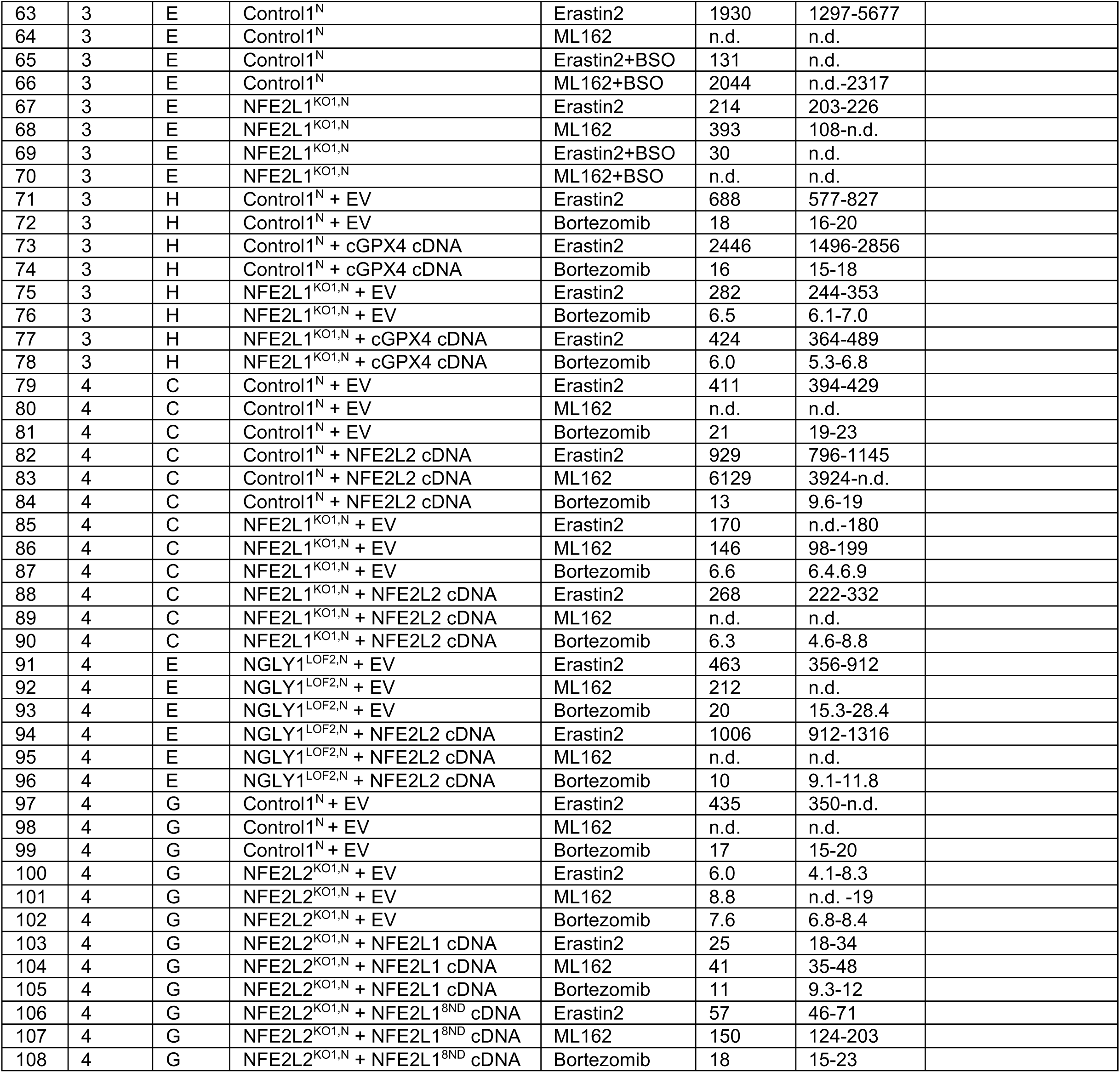
EC_50_ values for lethal treatment conditions. EV: empty vector, 95% C.I.: 95% confidence interval, n.d.: not determinable, n.s.: not significant. *Values appear in this table multiple times, as data from Figure 1*J*, 2*E*, and 2*I* were acquired within the same experiment.

## References

1. X. Jiang, B. R. Stockwell, M. Conrad, Ferroptosis: mechanisms, biology and role in disease. Nat Rev Mol Cell Biol 10.1038/s41580-020-00324-8 (2021).

2. W. S. Yang et al., Regulation of ferroptotic cancer cell death by GPX4. Cell 156, 317–331 (2014).

3. J. I. Leu, M. E. Murphy, D. L. George, Mechanistic basis for impaired ferroptosis in cells expressing the African-centric S47 variant of p53. Proc Natl Acad Sci U S A 116, 8390–8396 (2019).

4. M. A. Badgley et al., Cysteine depletion induces pancreatic tumor ferroptosis in mice. Science 368, 85–89 (2020).

5. S. J. Dixon et al., Ferroptosis: an iron-dependent form of nonapoptotic cell death. Cell 149, 1060–1072 (2012).

6. L. Magtanong et al., Exogenous Monounsaturated Fatty Acids Promote a Ferroptosis-Resistant Cell State. Cell Chem Biol 26, 420–432 e429 (2019).

7. G. P. Sykiotis, D. Bohmann, Stress-activated cap’n’collar transcription factors in aging and human disease. Sci Signal 3, re3 (2010).

8. N. Liu, X. Lin, C. Huang, Activation of the reverse transsulfuration pathway through NRF2/CBS confers erastin-induced ferroptosis resistance. Br J Cancer 122, 279–292 (2020).

9. N. Takahashi et al., 3D Culture Models with CRISPR Screens Reveal Hyperactive NRF2 as a Prerequisite for Spheroid Formation via Regulation of Proliferation and Ferroptosis. Mol Cell 80, 828–844 e826 (2020).

10. M. Dodson, R. Castro-Portuguez, D. D. Zhang, NRF2 plays a critical role in mitigating lipid peroxidation and ferroptosis. Redox Biol 23, 101107 (2019).

11. F. Kuang, J. Liu, Y. Xie, D. Tang, R. Kang, MGST1 is a redox-sensitive repressor of ferroptosis in pancreatic cancer cells. Cell Chem Biol 28, 765–775 e765 (2021).

12. Z. Fan et al., Nrf2-Keap1 pathway promotes cell proliferation and diminishes ferroptosis. Oncogenesis 6, e371 (2017).

13. X. Sun et al., Activation of the p62-Keap1-NRF2 pathway protects against ferroptosis in hepatocellular carcinoma cells. Hepatology 63, 173–184 (2016).

14. Y. P. Kang et al., Non-canonical Glutamate-Cysteine Ligase Activity Protects against Ferroptosis. Cell Metab 33, 174–189 e177 (2021).

15. S. Vomund, A. Schafer, M. J. Parnham, B. Brune, A. von Knethen, Nrf2, the Master Regulator of Anti-Oxidative Responses. Int J Mol Sci 18 (2017).

16. K. Chan, R. Lu, J. C. Chang, Y. W. Kan, NRF2, a member of the NFE2 family of transcription factors, is not essential for murine erythropoiesis, growth, and development. Proc Natl Acad Sci U S A 93, 13943–13948 (1996).

17. J. Y. Chan et al., Targeted disruption of the ubiquitous CNC-bZIP transcription factor, Nrf-1, results in anemia and embryonic lethality in mice. EMBO J 17, 1779–1787 (1998).

18. M. Kwong, Y. W. Kan, J. Y. Chan, The CNC basic leucine zipper factor, Nrf1, is essential for cell survival in response to oxidative stress-inducing agents. Role for Nrf1 in gamma-gcs(l) and gss expression in mouse fibroblasts. J Biol Chem 274, 37491–37498 (1999).

19. L. Chen et al., Nrf1 is critical for redox balance and survival of liver cells during development. Mol Cell Biol 23, 4673–4686 (2003).

20. L. Leung, M. Kwong, S. Hou, C. Lee, J. Y. Chan, Deficiency of the Nrf1 and Nrf2 transcription factors results in early embryonic lethality and severe oxidative stress. J Biol Chem 278, 48021–48029 (2003).

21. J. Steffen, M. Seeger, A. Koch, E. Kruger, Proteasomal degradation is transcriptionally controlled by TCF11 via an ERAD-dependent feedback loop. Mol Cell 40, 147–158 (2010).

22. S. K. Radhakrishnan et al., Transcription factor Nrf1 mediates the proteasome recovery pathway after proteasome inhibition in mammalian cells. Mol Cell 38, 17–28 (2010).

23. K. Itoh et al., Keap1 represses nuclear activation of antioxidant responsive elements by Nrf2 through binding to the amino-terminal Neh2 domain. Genes Dev 13, 76–86 (1999).

24. S. B. Cullinan, J. D. Gordan, J. Jin, J. W. Harper, J. A. Diehl, The Keap1-BTB protein is an adaptor that bridges Nrf2 to a Cul3-based E3 ligase: oxidative stress sensing by a Cul3-Keap1 ligase. Mol Cell Biol 24, 8477–8486 (2004).

25. A. L. Eggler, E. Small, M. Hannink, A. D. Mesecar, Cul3-mediated Nrf2 ubiquitination and antioxidant response element (ARE) activation are dependent on the partial molar volume at position 151 of Keap1. Biochem J 422, 171–180 (2009).

26. F. M. Tomlin et al., Inhibition of NGLY1 Inactivates the Transcription Factor Nrf1 and Potentiates Proteasome Inhibitor Cytotoxicity. ACS Cent Sci 3, 1143–1155 (2017).

27. A. B. Dirac-Svejstrup et al., DDI2 Is a Ubiquitin-Directed Endoprotease Responsible for Cleavage of Transcription Factor NRF1. Mol Cell 79, 332–341 e337 (2020).

28. N. J. Lehrbach, P. C. Breen, G. Ruvkun, Protein Sequence Editing of SKN-1A/Nrf1 by Peptide:N-Glycanase Controls Proteasome Gene Expression. Cell 177, 737–750 e715 (2019).

29. P. Lipari Pinto et al., NGLY1 deficiency-A rare congenital disorder of deglycosylation. JIMD Rep 53, 2–9 (2020).

30. A. Tsherniak et al., Defining a Cancer Dependency Map. Cell 170, 564–576 e516 (2017).

31. F. M. Behan et al., Prioritization of cancer therapeutic targets using CRISPR-Cas9 screens. Nature 568, 511–516 (2019).

32. J. Y. Cao et al., A Genome-wide Haploid Genetic Screen Identifies Regulators of Glutathione Abundance and Ferroptosis Sensitivity. Cell Rep 26, 1544–1556 e1548 (2019).

33. H. Dreger et al., Nrf2-dependent upregulation of antioxidative enzymes: a novel pathway for proteasome inhibitor-mediated cardioprotection. Cardiovasc Res 83, 354–361 (2009).

34. G. C. Forcina, M. Conlon, A. Wells, J. Y. Cao, S. J. Dixon, Systematic Quantification of Population Cell Death Kinetics in Mammalian Cells. Cell Syst 4, 600–610 e606 (2017).

35. S. C. Lu, Glutathione synthesis. Biochim Biophys Acta 1830, 3143–3153 (2013).

36. J. I. Leu, M. E. Murphy, D. L. George, Functional interplay among thiol-based redox signaling, metabolism, and ferroptosis unveiled by a genetic variant of TP53. Proc Natl Acad Sci U S A 117, 26804–26811 (2020).

37. A. Tarangelo et al., p53 Suppresses Metabolic Stress-Induced Ferroptosis in Cancer Cells. Cell Rep 22, 569–575 (2018).

38. L. Jiang et al., Ferroptosis as a p53-mediated activity during tumour suppression. Nature 520, 57–62 (2015).

39. W. H. Yang et al., The Hippo Pathway Effector TAZ Regulates Ferroptosis in Renal Cell Carcinoma. Cell Rep 28, 2501–2508 e2504 (2019).

40. J. Wu et al., Intercellular interaction dictates cancer cell ferroptosis via NF2-YAP signalling. Nature 572, 402–406 (2019).

41. Y. Yoshida et al., Loss of peptide:N-glycanase causes proteasome dysfunction mediated by a sugar-recognizing ubiquitin ligase. Proc Natl Acad Sci U S A 118 (2021).

42. F. Katsuoka, M. Yamamoto, Small Maf proteins (MafF, MafG, MafK): History, structure and function. Gene 586, 197–205 (2016).

43. A. Raghunath et al., Antioxidant response elements: Discovery, classes, regulation and potential applications. Redox Biol 17, 297–314 (2018).

44. J. Kong et al., Mitochondrial function requires NGLY1. Mitochondrion 38, 6–16 (2018).

45. K. Yang, R. Huang, H. Fujihira, T. Suzuki, N. Yan, N-glycanase NGLY1 regulates mitochondrial homeostasis and inflammation through NRF1. J Exp Med 215, 2600–2616 (2018).

46. S. Y. Han et al., A conserved role for AMP-activated protein kinase in NGLY1 deficiency. PLoS Genet 16, e1009258 (2020).

47. D. M. Talsness et al., A Drosophila screen identifies NKCC1 as a modifier of NGLY1 deficiency. Elife 9 (2020).

48. M. A. Tambe, B. G. Ng, H. H. Freeze, N-Glycanase 1 Transcriptionally Regulates Aquaporins Independent of Its Enzymatic Activity. Cell Rep 29, 4620–4631 e4624 (2019).

49. A. Galeone et al., Regulation of BMP4/Dpp retrotranslocation and signaling by deglycosylation. Elife 9 (2020).

50. K. G. Owings, J. B. Lowry, Y. Bi, M. Might, C. Y. Chow, Transcriptome and functional analysis in a Drosophila model of NGLY1 deficiency provides insight into therapeutic approaches. Hum Mol Genet 27, 1055–1066 (2018).

51. S. Iyer et al., Drug screens of NGLY1 deficiency in worm and fly models reveal catecholamine, NRF2 and anti-inflammatory-pathway activation as potential clinical approaches. Dis Model Mech 12 (2019).

52. F. A. Ran et al., Genome engineering using the CRISPR-Cas9 system. Nat Protoc 8, 2281–2308 (2013).

53. M. Conlon et al., A compendium of kinetic modulatory profiles identifies ferroptosis regulators. Nat Chem Biol 10.1038/s41589-021-00751-4 (2021).

